# Bovine blastocyst like structures derived from stem cell cultures

**DOI:** 10.1101/2022.12.20.521301

**Authors:** Carlos A. Pinzón-Arteaga, Yinjuan Wang, Yulei Wei, Leijie Li, Ana Elisa Ribeiro Orsi, Giovanna Scatolin, Lizhong Liu, Masahiro Sakurai, Jianfeng Ye, Leqian Yu, Bo Li, Zongliang Jiang, Jun Wu

**Affiliations:** Department of Molecular Biology, University of Texas Southwestern Medical Center, Dallas, TX, USA; School of Animal Sciences, AgCenter, Louisiana State University, Baton Rouge, LA, 70810, USA; State key laboratory of Agrobiotechnology, College of Biological Sciences, China, Agricultural University, Beijing, 100193, China; SJTU-Yale Joint Center for Biostatistics and Data Science, School of Life Sciences and Biotechnology, Shanghai Jiao Tong University, Shanghai, China; Department of Genetics and Evolutionary Biology, Institute of Biosciences, University of São Paulo, São Paulo, Brazil; Lyda Hill Department of Bioinformatics, University of Texas Southwestern Medical Center, Dallas, TX, USA; The State Key Laboratory of Stem Cell and Reproductive Biology, Institute of Zoology, Chinese Academy of Sciences, Beijing 100101, P. R. China; Institute for Stem Cell and Regeneration, Chinese Academy of Sciences, Beijing 100101, P. R. China; Department of Animal Sciences, Genetics Institute, University of Florida, Gainesville, Florida, 32610, USA; Hamon Center for Regenerative Science and Medicine, University of Texas Southwestern Medical Center, Dallas, TX 75390, USA; Cecil H. and Ida Green Center for Reproductive Biology Sciences, University of Texas Southwestern Medical Center, Dallas, TX, USA

**Author notes:** To whom correspondence will be addressed. These authors contribute equally (Carlos A. Pinzón-Arteaga, Yinjuan Wang, Yulei Wei).

## Abstract

Understanding blastocyst formation and implantation is critical for improving farm animal reproduction but is hampered by a limited supply of embryos. We developed an efficient method to generate bovine blastocyst-like structures (termed blastoids) via the assembly of trophoblast stem cells and expanded potential stem cells. Bovine blastoids resemble blastocysts in morphology, cell composition, single-cell transcriptomes, and represent an accessible in vitro model for studying bovine embryogenesis.

---

Blastoids, were initially developed in mice by assembling embryonic stem cells (ESCs)^1^ or extended pluripotent stem cells (EPSCs)^2^ with trophoblast stem cells (TSCs), or through EPSC differentiation and self-organization^3^, have also been successfully generated in humans^4-8^. To date, however, blastoids from other species have not been reported. Recently, several types of pluripotent stem cells (PSCs), including EPSCs, have been derived from Bos taurus blastocysts^9-15^, which hold great potential to advance animal agriculture^16^. Surprisingly, we found a bovine EPSC condition^13, 17^ could support de novo derivation and long-term culture of bovine trophoblast stem cells (TSCs) (Wang et al., manuscript co-submitted). The availability of bovine EPSCs and TSCs prompted us to test whether bovine blastoids could be generated through 3D assembly (**Extended Data Fig. 1a**).

To develop a condition that supports bovine blastoid formation, we adapted the FAC (<u>F</u>GF2, <u>A</u>ctivin-A and <u>C</u>HIR99021) medium^18^, as this medium supports the differentiation of hypoblast-like cells (HLCs) from naïve human PSCs^4^, and added the leukemia inhibitory factor (LIF) that is known to improve preimplantation bovine embryo development^19^ (FACL). FGF signaling level can bias the fate of inner cell mass (ICM) likely acting through the MEK-ERK pathway^20, 21^, where high level of FGF directs ICM cells towards the hypoblast (HYPO, or primitive endoderm [PE]) lineage ^22^. To support both HYPO and epiblast (EPI) lineages, we optimized FGF signaling by lowering FGF2 concentration and including a low dose of a MEK inhibitor (PD0325901, 0.3µM), as MEK inhibition has been shown to suppress HYPO fate in bovine embryos in a dose dependent manner^23^. This optimized condition, termed titrated FACL+PD03 (tFACL+PD) (see Methods), supported the formation of bovine blastoids with high efficiency (64.2±7.6%) within 4 days (**Fig. 1a, b, and Extended Data Fig. 1b-h)**. Morphologically each bovine blastoid contains a cavity, an outer trophectoderm (TE)-like layer and an ICM-like compartment, which resembles bovine blastocysts produced by *in vitro* fertilization (IVF) (**Fig. 1b, and Supplementary Video 1)**. Blastocele and ICM sizes of day-4 bovine blastoids reached diameters equivalent to day-8 IVF blastocysts (**Fig. 1c, d)**. We performed immunofluorescence (IF) analysis and found bovine blastoids expressed markers characteristic of EPI (SOX2), HYPO (SOX17) and trophectoderm (TE) (GATA3, KRT18, and CDX2) lineages, and stained positive for a tight junction marker ZO1(TJP1) and an apical marker F-actin (Phallodin) (**Fig. 1e, Extended Data Fig. 2**). Despite the similarities, we found that the expression levels of some lineage markers were different between blastocysts and blastoids when quantified via IF, with blastoid trophoblast-like cells (TLCs) expressing higher levels of CDX2 and HLCs, and epiblast-like cells (ELCs) expressing lower levels of SOX17 and SOX2 when compared to their corresponding cell types in blastocysts produced by *in vitro* fertilization (IVF) (**Extended Data Fig. 2e**). Analysis of lineage composition in bovine blastoids by flow cytometry further revealed that on average 49.67 ±3.29%, 31.47±2.54%, 6.58 ±1.85% cells stained positive for SOX2 (EPI), CDX2 (TE), and SOX17 (HYPO), respectively (**Fig. 1f, Extended Data Fig. 3**).

**Figure 1.**
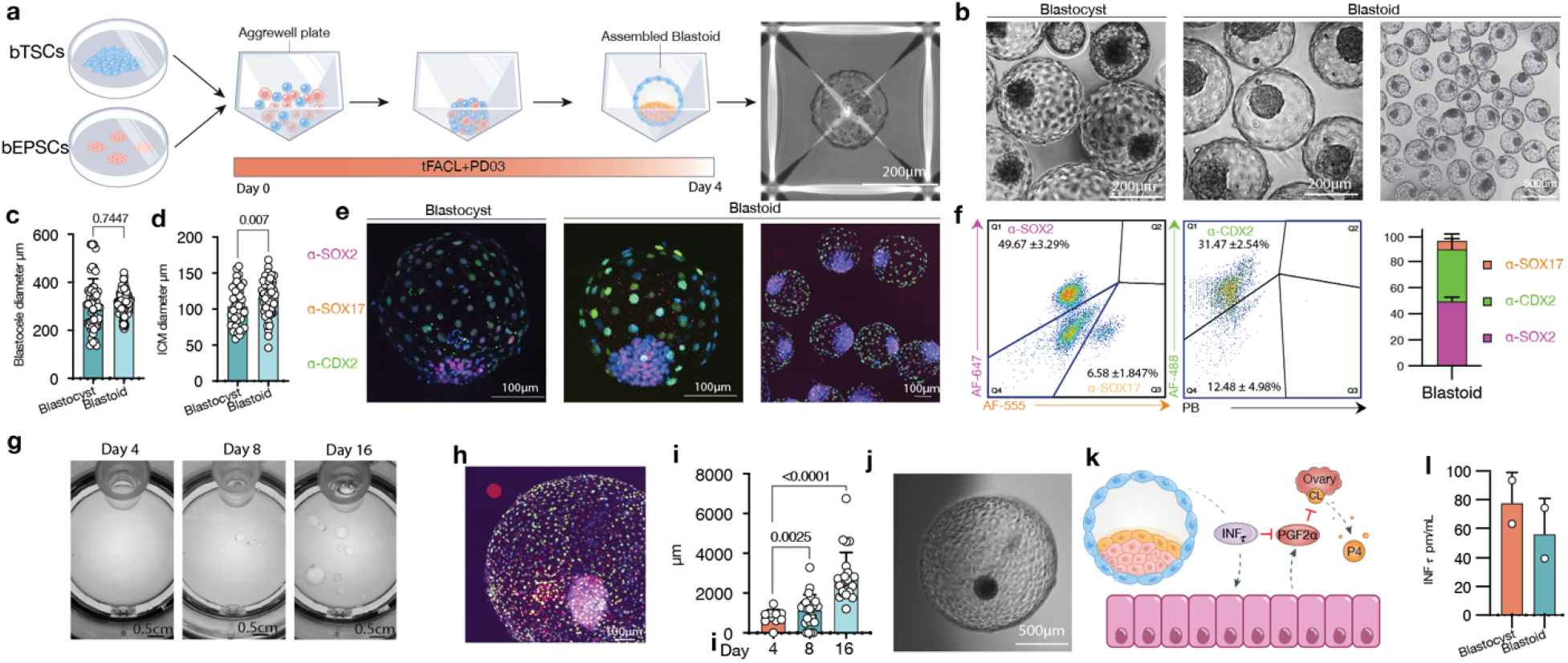
Assembly of bovine blastoids from EPSCs and TSCs cultures. **a**. Illustration of the assembly process via bovine EPSCs and TSCs aggregation. **b**. Phase-contrast image comparing blastoids vs blastocysts. **c**. Blastocele diameter measurement. **d**. Inner cell mass (ICM) diameter measurement. **e**. Immunostaining for epiblast marker SOX2 (magenta, EPI), hypoblast marker SOX17(red, HYPO) and trophectoderm marker CDX2(green, TS), individual markers in Extended Data 1 and 2. **f**. Flow cytometry quantification of single cell dissociated blastoids showing the relative quantities for each lineage, left panel cells are gated from SOX2 and SOX17 negative (Q4) cells in right panel, n=3. **g**. Snapshots of in vitro growth of blastoids in rotating culture system (Clinostar Incubator, Celvivo). **h**. Representative image via immunostaining of all three lineages as in e, individual markers in Extended Data figure 4. **i**. Blastoid diameter quantification. **j**. representative micrographs of in vitro grown blastoid. **k**. A schematic of the maternal recognition of the action of pregnancy signal interferon TAU (INFt). **l**. Enzyme-linked immunosorbent assay (ELISA) measurement of (INFt) in surrogate recipients following embryo transfers. PGF2α: Prostaglandin F2α. P4: Progesterone.

Next, we evaluated the *in vitro* growth of blastoids and blastocysts under a 3D rotating culture (see Methods). We found trophoblast cells and cavities in both IVF blastocysts and blastoids continued to proliferate and expand over a period of more than 2 weeks, which were also accompanied by an increase in the ICM size (**Fig. 1g-j, Extended Data Fig. 4, and Supplementary Video 2**). To evaluate whether blastoids can establish pregnancy, we performed embryo transfer to synchronized surrogates (see Methods). Interestingly, we were able to detect the anti-luteolytic hormone interferon-tau (INF*τ*) in the surrogate blood. INF*τ* is the signal for maternal recognition of pregnancy in ruminants, which acts by blocking prostaglandin release from the uterus and allowing the corpus luteum to persist and the pregnancy to be maintained^24-26^ (**Fig. 1k)**. INF*τ* was measured at concentrations of 56.53±25.13pm/ml in 2 out of 4 surrogates 7 days following blastoid transfer, which were comparable to those from IVF blastocyst transfers (78.36±21.54pm/ml) in 2 out of 5 surrogates (**Fig. 1l**).

To determine the transcriptional states of bovine blastoid cells, we performed single-cell RNA-sequencing (scRNA-seq) using the 10x Genomics Chromium platform and carried out integrated analysis with Smart-seq2 single-cell transcriptomes derived from zygote^27^, 2-cell^27^, 8-cell^28^, 16-cell^28^, morula^27^ and day 7.5 blastocyst stage IVF bovine embryos^27^ as well as *in vivo* bovine blastocysts (see data availability). Joint uniform manifold approximation and projection (UMAP) embedding revealed blastoid-derived cells clustered with blastocyst-derived cells (**Fig. 2a, b**). To further evaluate the temporal identity of blastoid cells, we performed pseudo bulk analysis on the 10x blastoid data to compensate for the differences in sequencing depth to Smart-seq2 data. For this analysis we also included datasets from bovine early gastrulation-stage embryos ^29^. We found that different embryo datasets were orderly arranged on the PCA plot according to their developmental time and blastoid cells were mapped closer to blastocyst cells **(Fig. 2c, d, and Extended Data Fig. 5**).

**Figure 2.**
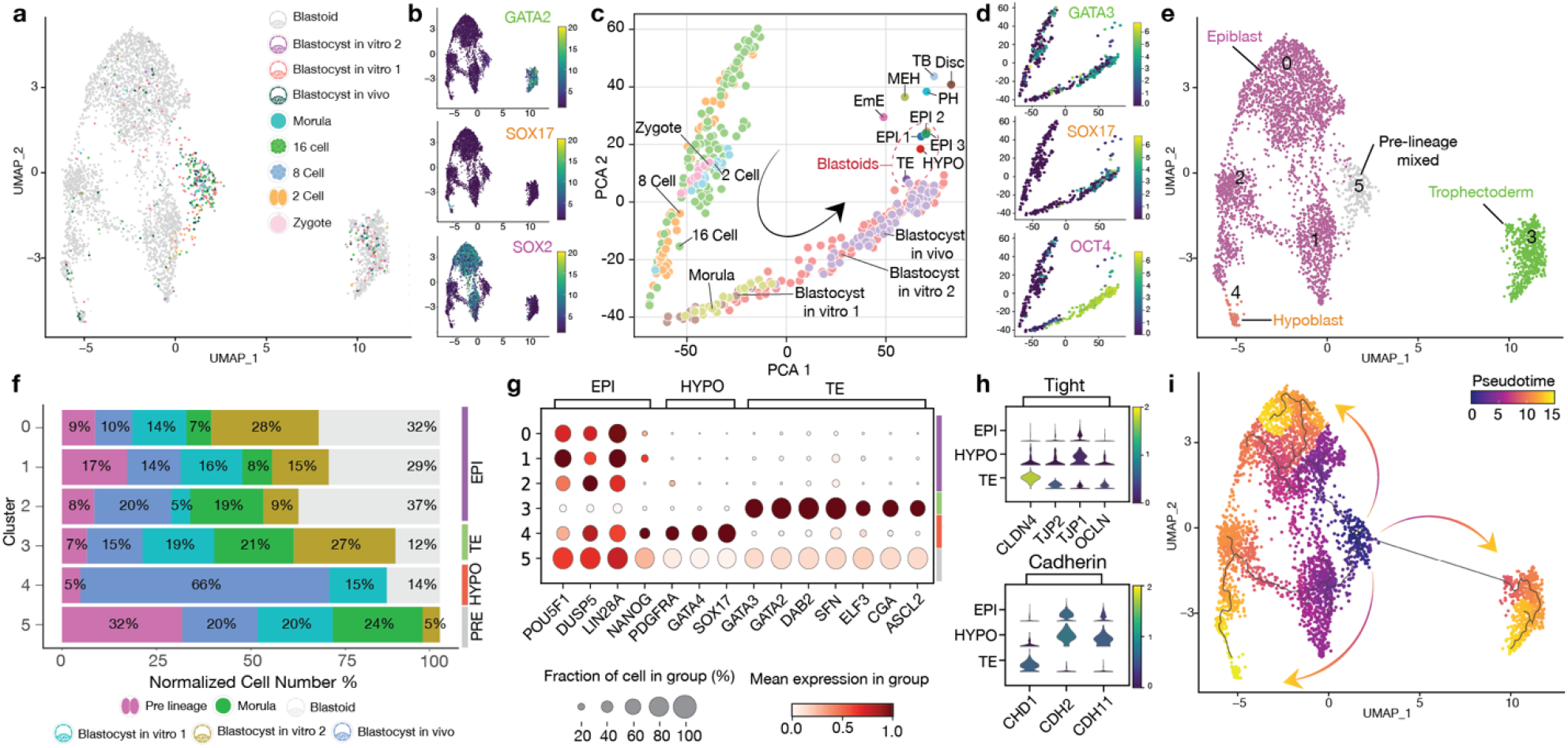
Single cell characterization of bovine assembled blastoids. **a**. Joint uniform manifold approximation and projection (UMAP) embedding of 10x Genomics single-cell transcriptomes of bovine blastoids (grey) and bovine zygote (pink), 2 cell (orange), 8 cell (blue), 16 cell (green), Morula (cyan) and *in vivo* and *in vitro* Blastocyst stage embryos (purple, dark green, light red). **b**. UMAP Heatmap showing expression of Trophectoderm (TE), Hypoblast (HYPO), and epiblast (EPI) markers, GATA2, SOX17 and SOX2, respectively **c**. Principal component analysis (PCA) of pseudo bulk conversion of blastoid data. Gastrulation markers^29^: Disc: Embryonic disc (Day 14 Stage 4). EmE: Embryonic ectoderm (Day 14 Stage 5). MEH: Mesoderm, endoderm and visceral hypoblast. (Day 14 Stage 5). PH: Parietal hypoblast. (Day 14 Stage 5). TB: Trophoblast. (Day 14 Stage 5). **d**. PCA heatmaps showing expression of Trophectoderm (TE), Hypoblast (HYPO), and epiblast (EPI) markers, GATA3, SOX17 and OCT4 (also known as POU5F1), respectively. **e**. Major cluster classification based on marker expression. **f**. Normalized percentage of cells in each cluster. **g**. Dot plot indicating the expression of markers of epiblast (EPI), trophectoderm (TE) and hypoblast (HYPO). **h**. Violin plot of lineage specific cell junction markers. **i**. RNA velocity pseudotime analysis depicting the cell trajectories.

We annotated the six identified cell clusters based on marker gene expression and overlap with cells from bovine embryos (**Fig. 2e-g**). Cluster 3 expresses TE markers, e.g., GATA2 and GATA3, and is annotated as TLCs; Cluster 4 expresses HYPO markers, e.g., GATA4 and SOX17, and thus represents HLCs; Three clusters (0, 1, 2) express EPI markers, e.g., SOX2 and LIN28a, and are designated as ELCs; Cluster 5 is mostly composed of cells from pre-blastocyst stage embryos (named pre-lineage), and each blastoid cluster expressed lineage specific cadherin and tight junction markers **(Fig. 2e-h, Extended Data Fig. 6**). To evaluate the relationship between clusters, we performed pseudo time analysis, which predicted the differentiation trajectories from pre-lineage cluster to blastocyst and blastoid lineages (**Fig. 2i, Extended Data Fig. 7**). Finally, cross-species comparison revealed similarities and differences of bovine blastoids with human blastoids and blastocysts (**Extended Data Fig. 8**).

In sum, here we report an efficient and robust protocol to generate bovine blastoids by assembling EPSCs and TSCs that can self-organize and faithfully recreate all blastocyst lineages. The bovine blastoids show resemblance to bovine blastocysts in morphology, size, cell number, lineage composition and allocation, and could produce maternal recognition signal upon transfer to recipient cows. The bovine blastoids represent a valuable model to study early embryo development and understand the causes of early embryonic loss. Upon further optimization, bovine blastoid technology could lead to the development of new artificial reproductive technologies for cattle breeding, which may enable a paradigm shift in livestock reproduction.

## Supporting information

Supplementary Video 1. Bovine blastoid Z plane overview.

Supplementary Video 2. Bovine blastoid 3D rotating culture on day 16.

## Materials and Methods

### Bovine EPSCs stem cell culture

Bovine female ESCs cultured in NBFR^10^ were adapted to the bEPSC^30^ condition via culture adaptation for a minimum of 5 passages, until a doom morphology was visible. Cell cultures were performed in 0.1% gelatin-coated 6 well plates with 5×10^5^ irradiated MEF / STO per well. Upon passaging cells were washed with 1xPBS and dissociated with TrypLE (Thermo Fisher) for 3 minutes at 37°C, cells were then collected with 0.05% BSA in DMEM-F12 (Thermo Fisher) and centrifuged at 1000xg for 3 minutes and resuspended in 1ml of media per 9.6cm^2^. Each passage cells were count using Countess II (Thermo Fisher) and plated at a density of 30,000cells/cm^2^, at this plating ratio cells were passaged every 4 days. Upon plating cells were treated with the CETP cocktail^31^, 50 nM chroman-1 (C, Tochris), 5 μM emricasan (E, Selleckchem), 0.7 μM trans-ISRIB (T, Tochris), and polyamine supplement (P, Thermo) diluted 1x (CETP) was routinely used during the first 12h after passaging. Fresh culture media was added every day. Cells were cultured at 37°C in a 5% CO_2_ humidified incubator. Cells were cryopreserved in bEPSC media with 10% DMSO at 0.5×10^6 cells per ml. Detailed description of medias are in Supplementary Table 1.

### Bovine TSCs stem cell culture

TSCs were cultured in LCDM media (Wang et al., manuscript co-submitted) as stated above with slight modifications (Supplementary Table 1). Upon passaging cells media was removed and treated with Accumax (Thermo Fisher) for 5 minutes at 37°C (No 1xPBS wash), cells were collected with the same volume of bTSC medium and gently lifted of the plate using a wide opening p100 pipette tip and gentle force. Cells were split in a 1:3 ratio and plated in mouse feeder cells with CETP. Only one ml of media was plated in a 6 well for the first 24h to facilitate TSCs attachment. TSCs do not survive well single cell dissociation and tend to form trophospheres if not plated correctly. These steps are critical for the continuous culture and expansion of these cells. Cells were incubated at 37°C in a 5% CO_2_ humidified incubator. Cells were cryopreserved in CoolCell lx cell freezing vial containers (Corning) in 45 % LCDM 45% FBS and 10% DMSO orProFreeze Freezing medium (Lonza, 12-769E) at 2×10^6 cells per ml.

### Blastoid formation

For blastoid formation EPSCs cells were collected as stated above. Bovine TSCs were washed with 1x PBS, dissociated with Trypsin for 10 minutes at 37 °C, and inactivated with DMEM-F12 containing 10% fetal bovine serum (FBS). Cells were washed twice and on final resuspension in their normal culture media with 1x CETP and 10 UI per ml of DNase I (Thermo Fisher). To deplete MEF, cells were placed in precoated 12 well plates (Corning) with 0.1% gelatin and incubated for 15 minutes at 37°C. Single cell dissociation was made by gentle but constant pipetting and by passing the cells through a glass capillary attached to a p200 pipette tip, pulled to an inner diameter of 40-60µm (micropipette puller, Sutter Instruments). After this, cells were collected and strained using a 70µm and then a 37µm cell strainers (Corning). This same single cell dissociation procedure was used for blastoids processing. Cells were stained with 1x trypan blue and manually counted in a Neubauer chamber. Current protocol is optimized for 16 bEPSCs and 16 bTSCs per well in a ∼1200 well Aggrewell 400 microwell culture plate (Stemcell technologies) for a total of 19,200 of each cell types per well. Each well was precoated with 500µl of Anti-Adherence Rinsing Solution (Stemcell technologies) and spun for 5 minutes at 1500 rcf. Wells were rinsed with 1ml of PBS just before aggregation. An appropriate number of cells for the wells to be aggregated were centrifuged at 1000xg for 3 minutes and resuspended in 1ml of tFACL + 0.3µM PD03 media per well, supplemented with 1x CETP. To ensure even distribution, each microwell was gently mixed by pipetting with a P200 pipette to ensure equal distribution of the cells along the microwell, then the plate was centrifuged at 1300xg for 2 minutes and put in a humidified incubator at 37°C with 5% CO_2_ and 5% Oxygen. As MEK inhibition inhibits hypoblast differentiation a gradual decrease can be done if higher numbers of hypoblast cells are desired from 0.3 to 0.125µM.

### In vitro fertilization

Bovine IVF was performed as previously described ^32^ with modifications. Briefly oocytes were collected at a commercial abattoir (DeSoto Biosciences) and shipped in an MOFA metal bead incubator (MOFA Global) at 38.5°C overnight in sealed sterile vials containing 5% CO_2_ in air-equilibrated Medium 199 with Earle’s salts (Thermo Fisher), supplemented with 10% fetal bovine serum (Hyclone), 1% penicillin–streptomycin (Invitrogen), 0.2-mM sodium pyruvate, 2-mM L-glutamine (Sigma), and 5.0 mg/mL of Folltropin (Vetoquinol). The oocytes were matured in this medium for 22 to 24 hours. Matured oocytes were washed twice in warm Tyrode lactate (TL) HEPES supplemented with 50 mg/mL of gentamicin (Invitrogen) while being handled on a stereomicroscope (Nikon) equipped with a 38.5°C stage warmer. In vitro fertilization was conducted using a 2-hour pre-equilibrated IVF medium modified TL medium supplemented with 250-mM sodium pyruvate, 1% penicillin–streptomycin, 6 mg/mL of fatty acid–free BSA (Sigma), 20-mM penicillamine, 10-mM hypotaurine, and 10 mg/mL of heparin (Sigma) at 38.5 C, 5% CO_2_ in a humidified air incubator. Frozen semen (Bovine-elite) was thawed at 35°C for 1 minute, then separated by centrifugation at 200xg for 20 minutes in a density gradient medium (Isolate, Irvine Scientific) 50% upper and 90% lower. Supernatant was removed; sperm pellet was resuspended in 2-mL modified Tyrode’s medium and centrifuged at 200 g for 10 minutes to wash. The sperm pellet was removed and placed into a warm 0.65-mL microtube before bulk fertilizing in Nunc four-well multidishes (VWR) containing up to 50 matured oocytes per well at a concentration of 1.0×10^6 sperm/mL. 18 hours after insemination, oocytes were cleaned of cumulus cells by constant pipetting for 3-minutes in vortex in 100µl drop of TL HEPES with 0.05% Hyaluronidase (Sigma), washed in TL HEPES, and then cultured in 500µl of IVC media (IVF-Biosciences) supplemented with 0.5xN2B27 (Thermo Fisher) and FLI^19^ under mineral oil (Irvine Scientific) cultured until the blastocyst stage. Cleavage rates were recorded on Day 2, and viable embryos were separated from nonviable embryos. Blastocyst rates were recorded on Day 8 after IVF.

### Immunofluorescent staining

Samples (Cells, single cells, blastoids and blastocysts) were fixed with 4% paraformaldehyde (PFA) in 1xDPBS for 20 min at room temperature, washed in wash buffer (0.1% Triton X-100, 5% BSA in 1xDPBS) for 15 minutes and permeabilized with 1% Triton X-100 in PBS for 1 h. For phosphor antibodies samples were treated with 0.5% SDS for 1h. Samples were then blocked with blocking buffer (PBS containing 5% Donkey serum, 5% BSA, and 0.1% Triton X-100) at room temperature for 1 h, or overnight at 4 °C. Because of the large number of blastoids, to facilitate processing blastoids were gently washed out of the aggrewell plate and separated from cell debris using a 100µM reversible strainer (Stem cells), blastoids were then placed in a 70µm strainer (Corning) in a 6 well plate containing wash buffer and the strainer was moved from one well to another between steps. Primary antibodies were diluted in blocking buffer according to supplementary table 1. Blastoids were incubated in primary antibodies in 96 wells for 2 h at room temperature or overnight at 4 °C. Samples were washed three times for 15 minutes with wash buffer, and incubated with fluorescent-dye conjugated secondary antibodies (AF-488, AF-555 or AF-647, Invitrogen) diluted in blocking buffer (1:300 dilution) for 2 h at room temperature or overnight at 4 °C. Samples were washed three times with PBS-T. Finally, cells were counterstained with 300 nM 4′,6-diamidino-2-phenylindole (DAPI) solution at room temperature for 20 min. Phalloidin was directly stained along with other secondary antibodies in blocking buffer.

### Imaging

Phase contrast images were taken using a hybrid microscope (Echo Laboratories, CA) equipped with objective x2/0.06 numerical aperture (NA) air, x4/0.13 NA air, x10/0.7 NA air and 20x/0.05 NA air. Fluorescence imaging was performed on 8 well µ-siles (Ibidi) on a Nikon CSU-W1 spinning-disk super resolution by optical pixel reassignment (SoRa) confocal microscope with objectives x4/0.13 NA, a working distance (WD) of 17.1nm, air; ×20/0.45 NA, WD 8.9–6.9 nm, air; ×40/0.6 NA, WD 3.6–2.85 nm, air.

### Imaging analysis

Imaging experiments were repeated at least twice, with consistent results. In the figure captions n denotes the number of biological repeats. Raw images were first processed in Fiji^33^ to create maximal intensity projection (MIP) and an export of representative images. Nuclear segmentation was performed in Ilastik. MIP images and segmentation masks were processed in MATLAB (R2022a) using custom code, which is available in a public repository. Nuclear localized fluorescence intensity was computed for each cell in each field, and the value was then normalized to the DAPI intensity of the same cell. Intensity values of all cells were plotted as mean ± s.d. Total cells and CDX2, SOX2 and SOX17 positive cell numbers were calculated with Imaris (v.9.9, Oxford).

### Flow Cytometry

Blastoids were collected under a stereo microscope and single cell dissociated as stated above for the TSCs. Strained single cells were processed as stated above for immunofluorescent staining performing wash steps in 1.5ml Eppendorf tubes on a 90° centrifuge. Flow cytometry was performed using the appropriate unstained and single stain controls in a DBiosciences LSR II flow cytometer and analyzed using Flow Jo. Gating Strategy is shown in Extended Data figure 3.

### In vitro growth

Prior to use for bovine blastoid culture, the water beads, inside the humidity chamber of the ClinoReactor (CelVivo), were hydrated with sterile water (Corning) overnight at 4°C. Once hydrated and the growth chamber was filled with N2B27 basal media, and the reactor chamber was equilibrated for 1h at 37°C before exchanging for culture media. For rotating-culture blastoids were collected at day 4 post aggregation and placed in pre-equilibrated ClinoReactors in 10ml of tFACL+PD03 media and 1x CETP (Supplementary table 1). ClinoReactors were placed in the ClinoStar incubator at 37 °C with a gas mix of 5%CO_2_, 5% O_2_ and air. The rotation speed was set between 10 and 12 rpm and was lowered progressively as the blastoids expanded. Optimal growth conditions were achieved by exchanging media every four days. Blastoid and blastocysts growth was also tested on N2B27 with rock inhibitor (Y27632) and activin A as reported in ^34^. (Extended Data Figure 1 h, I)

### Embryo Transfer

Surrogate cows were synchronized with an intramuscular (IM) an injection of ovulation-inducing gonadotropin-release hormone (GnRH, Fertagyl), followed by a standard 7-day vaginal controlled drug internal release (CIDR) of progesterone. Upon CIDR removal, one dose of prostaglandin (Lutaluse) was administered. 48 hours after CIDR removal another dose of GnRH was administered via IM injection. A cohort of 15-20 bovine blastoids or 12-15 control IVF blastocysts were loaded into 0.5 mL straws in prewarmed Holding medium (ViGro) and transferred non-surgically to the uterine horn ipsilateral to the ovary with the corpus luteum (CL) as detected by transrectal ultrasound. 7 days after transfer, blastoids where be recovered by standard non-surgical flush with lactated ringers’ solution supplemented with 1% fetal bovine serum. All recipients were treated with prostaglandin (Lutaluse) after flushing.

### Quantitative measurement of Bovine IFN-*tau* in blood

Blood samples from surrogate and controls were drawn from the coccygeal vein using serum separator tubes. The samples were immediately placed in refrigerator overnight before centrifugation for 15 minutes at 1000 ×g. IFN***τ*** in the serum was determined by Bovine Interferon-Tau ELISA Kit (CSB-E 16948B) according to manufacturer’s protocol. Briefly, each well was added 100 µL standard or sample and incubated for 2 hours at 37 □. Then, liquid was removed and 100 µL Biotin-antibody (1X) was added to each well, incubating 1 hour at 37 □. After aspirating the wells, 200 µL Wash Buffer was used to wash the wells for three times. After last wash, the plate was inverted and blotted against clean paper towels to remove any remaining Wash Buffer. 100 µL HRP-avidin (1X) was added to each well and incubated for 1 hour at 37 □. 200 µL Wash Buffer was used to wash the wells for five times. 90 µL TMB Substrate was added and incubated for 20 minutes at 37 □. Protect from light. 50 µL Stop Solution was added to each well, gently tapping plate to ensure thorough mixing. The plate was measured using microplate reader set to 450 nm.

### Single-cell RNA-Seq library generation

Bovine blastoids were single cell dissociated and strained cells were prepared as stated adobe. Cells were washed in PBS containing 0.04% BSA and centrifuged at 90° x500g for 5 min. Cell were resuspended in PBS containing 0.04% BSA at a single cell suspension of 1,000 cells/µL. Cells were loaded into a 10x Genomics Chromium Chip following manufacturer instruction (10x Genomics, Pleasanton, CA, Chromium Next GEM Single Cell 3L GEM, Library & Gel Bead Kit v3.1) and sequenced by Illumina NextSeq 500/550 sequencing systems (Illumina).

### Published single-cell data collection

We collected single-cell sequencing data from published literature for comparative analysis. Two Bovine IVF single-cell sequencing raw FASTQ data were downloaded from the GEO database, including 179 IVF cells^35^ sequenced using Smart-seq2 and 98 IVF cells^36^ sequenced using STRT-seq.

### Pre-processing single-cell data

For 10X Genomics single-cell data, we used the Cell Ranger pipeline (v.3.1.0) with default parameters to generate the expression count matrix. The bovine reference genome and gene annotation file were downloaded from Ensembl database (UMD3.1) and generated by Cell Ranger mkfastq with default parameters. Seurat^37^ (3.1.4) was used to single-cell quality control. To reduce multiplets and dead cells, we screened cells with expressed gene numbers between 200 and 6000, unique molecular identifiers (UMIs) between 5000 and 30,000, and mitochondrial RNA genes counts below 15 percent.

For public Smart-seq2 and STRT-seq data, raw FASTQ reads were trimmed using Trim Galore (0.6.4, https://www.bioinformatics.babraham.ac.uk/projects/trim_galore/) with default parameters. In order to minimize processing differences, trimmed reads were aligned to the same genome reference (UMD3.1) by using HISAT2^38^ (2.1.0) with default parameters. Read counts per gene were annotated by HTSeq-count^39^ software (2.0.2) using the same gene annotation files (UMD3.1). Then, transcripts per million (TPM) were calculated to reduce gene length differences. Also, dead cells were removed by filter mitochondrial gene counts content below 15%.

### Normalization and dimensionality reduction

We used log-percentage value to normalize each single-cell expression matrix, which can reduce the bias of gene expression values caused by different sequencing depths and sequencing methods. In order to reducing the dimension of feature genes and improving the efficiency and accuracy of integration, the variance and mean of genes in each single-cell cohort were used to fit local polynomial regression and filter the top 2000 variable feature genes^40^.

### Data integration and clustering

The Find Integration Anchors model in the Seurat package was used to find the similarity anchor structure between different single-cell data. Then, we completed the data integration according to the anchors information with 80 dimensions, 20 anchors, 40 candidate cells, and reciprocal PCA for dimensionality reduction (‘dims = 1:80, k.anchor = 20, k.filter = 40, reduction = “rpca”‘). Single cells were clustered using the shared nearest neighbor (SNN) modularity optimization-based clustering algorithm in Seurat package, with 90 Principal Component (PC) and 0.6 resolution. Then, Uniform manifold approximation and projection (UMAP) was used to reduce the dimensions and show the visualize figure with non-default parameters: ‘dims = 1:90’.

### Gene function annotation

Gene ontology (GO)^41^ terms and Kyoto encyclopedia of genes and genomes (KEGG)^42^ pathways enrichment were performed using clusterProfiler^43^ (3.14.3; org.Bt.eg.db v 3.10.0) with parameter: ‘pvalueCutoff = 0.05’.

### Pseudotime construction

Monocle3^44^ (0.2.3.0) was used for pseudotime analysis, with the UMI matrix and UMAP embedding matrix generated by Seurat as input. Cell pseudotime trend was learnt by using cells in all clusters to generate a single and acyclic structure graph (‘use_partition = F, close_loop = F’).

## Data availability

8 cell and 16 cell from GSE99210 (Single-cell RNA sequencing reveals developmental heterogeneity of blastomeres during major genome activation in bovine embryos)^28^. Zygote, 2 cell, 8 cell, morula and blastocyst from PRJNA727165 (Reprogramming barriers in bovine cells nuclear transfer revealed by single□cell RNA□seq analysis)^27^. Raw unprocessed data of gastrulation embryos was obtained from Dr. Peter L. Pfeffer^29^. *In vivo* blastocyst and *in vitro* blastocyst1 datasets were obtained from Dr. Zongliang Jiang (GSE215409). Bovine blastoid single cell raw and processed data have been deposited in the Gene Expression Omnibus under accession code (GSE221248).

## Author contributions

C.A.P-A., Y.Wang., Z.J. and J.W. conceptualized, designed, analyzed, and interpreted the experimental results. C.A.P-A., Y.Wang. and Y.Wei. performed blastoid generation experiments. M.S. helped with in vitro fertilization of bovine embryos. Y.W. and L.Y. helped with immunostaining. C.A.P-A. and Y.Wang. performed extended in vitro culture of bovine blastocysts and blastoids. C.A.P-A., G.S., Y.Wang. and Z.J. performed embryo transfer experiments. J.Y. and B.L. prepared scRNA-seq library. C.A.P-A., L.I. and A.E.R.O. performed scRNA-seq analysis. Z.J. and J.W. supervised the study. C.A.P-A., Z.J. and J.W. wrote the manuscript with inputs from all authors.

## Acknowledgements

We thank Dr. Joel Carter from J A Carter, CETS, LLC, for his assistance with embryo transfer. J.W. is a New York Stem Cell Foundation–Robertson Investigator and Virginia Murchison Linthicum Scholar in Medical Research and funded by CPRIT (RR170076), NIH (GM138565-01A1 and OD028763), Welch (854671) and UT Southwestern & Texas A&M clinical translation and translational award (CSTA) program supported by the NIH (1UL1TR003163-01A1). Z.J. is funded by the United States Department of Agriculture (2019–67016–29863), the National Institutes of Health (R01HD102533). The Nikon SoRa spinning disk microscope was purchased by the UTSW quantitative light microscopy core with a Shared Instrumentation grant from NIH award: 1S10OD028630-01 to Katherine Luby-Phelps.

## Conflict of interests

C.A.P-A., Y.Wang., Y.Wei., Z.J. and J.W. are co-inventors on US provisional patent application 63/370,068 relating to the Bovine blastocysts like structures and uses thereof.

**Supplementary Table 1**.

1. Culture media

2. Material lists

3. Antibodies

**Supplementary Video 1. Bovine blastoid Z plane overview**.

**Supplementary Video 2. Bovine blastoid 3D rotating culture on day 16**.

**Extended Data figure 1.**
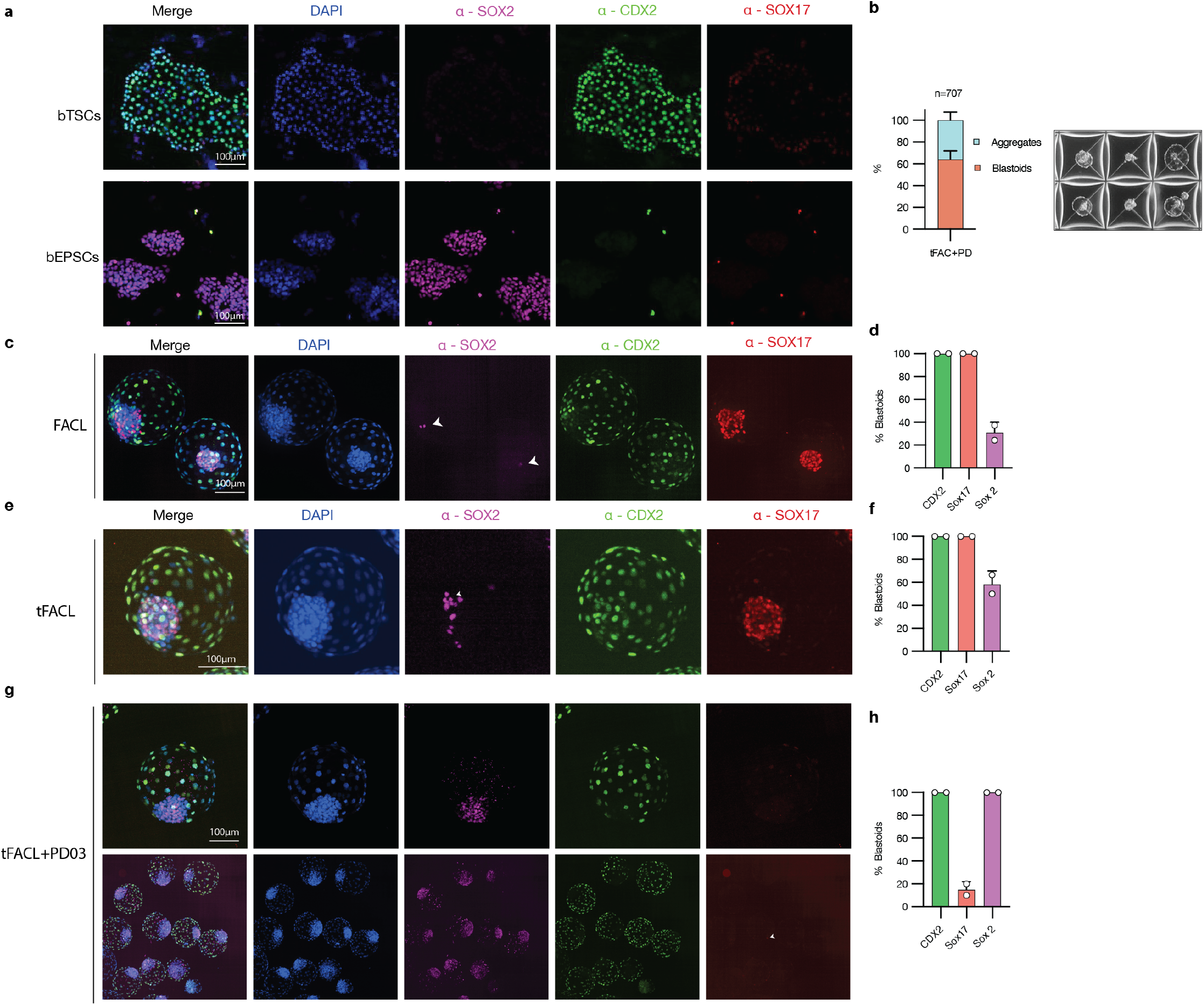
Stem cell cultures and Blastoid media optimization. **a**. Immunostaining of bovine EPSCs and TSCs for epiblast marker SOX2 (cyan), hypoblast marker SOX17(red) and trophectoderm marker CDX2 (green). **b**. Quantification of blastoid formation efficiency. Immunostaining for epiblast marker SOX2 (magenta), hypoblast marker SOX17(red) and trophectoderm marker CDX2(green) and marker quantification **c-d**. FACL. **e-f**. tFACL. **g-h**. FACL+ PD.

**Extended Data figure 2.**
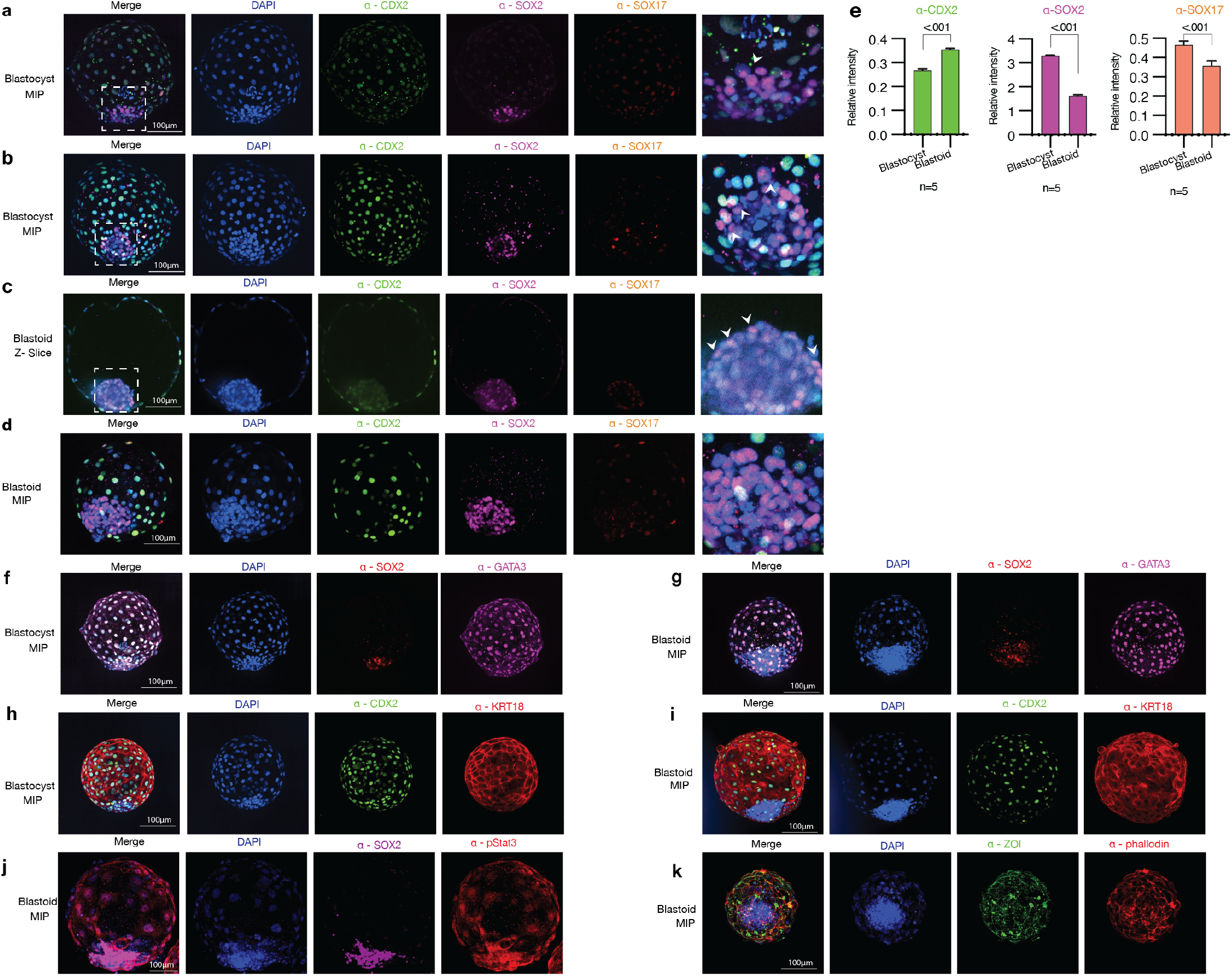
Bovine blastoids immunostaining characterization and comparison to IVF blastocysts. Immunostaining for epiblast marker SOX2 (magenta), hypoblast marker SOX17(red) and trophectoderm marker CDX2(green) **a-b**. IVF Blastocysts and **c-d**. Blastoids. **e**. DAPI normalized relative intensity quantification of side-by-side staining and imaging of blastocysts and blastoids n=5, mean ± s.d. Immunostaining for epiblast marker SOX2 (red), and trophectoderm marker gata3(magenta). **f**. Blastocyst and **g**. Blastoid. Immunostaining for trophectoderm markers CDX2(green) and Keratin 18 (red). **f**. Blastocyst and **g**. Blastoid. J. Immunostaining for phospho-STAT3(red). **k**. Immunostaining for tight junction marker ZO1(TJP1, green) and apical marker F-actin (Phallodin, Red).

**Extended Data figure 3.**
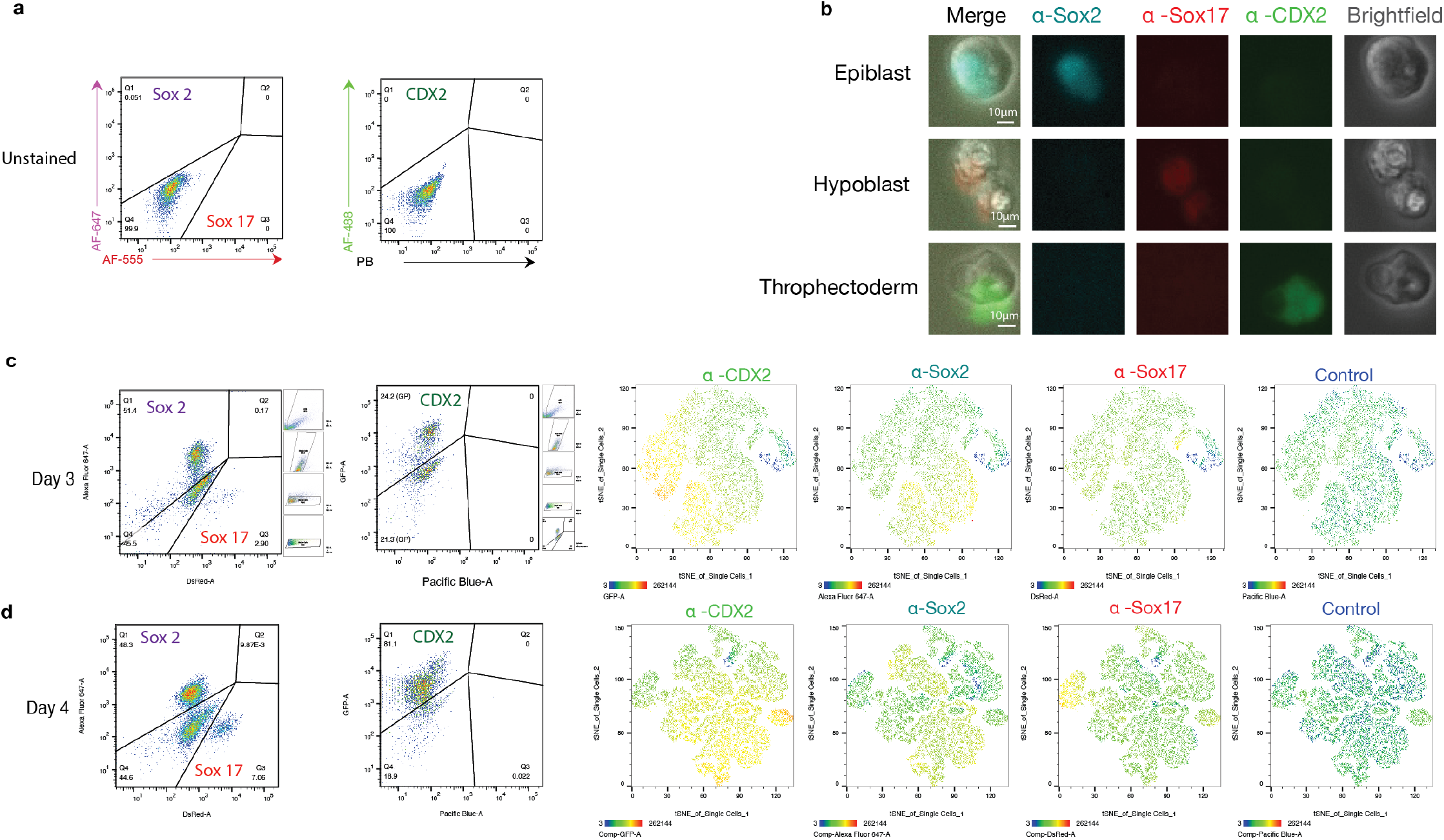
Blastoids lineage quantification by flow cytometry. **a**. Unstained control. **b**. Imaging examples of stained cells quantified by flow cytometry. **c-d**. Lineage quantification for epiblast marker SOX2 (AF-647), hypoblast marker SOX17(AF-555, DsRed channel) and trophectoderm marker CDX2(AF-488, GFP channel), autofluorescence control (Pacific blue) and tSNE plots of each of the quantified markers for days 3 and 4 of protocol.

**Extended Data figure 4.**
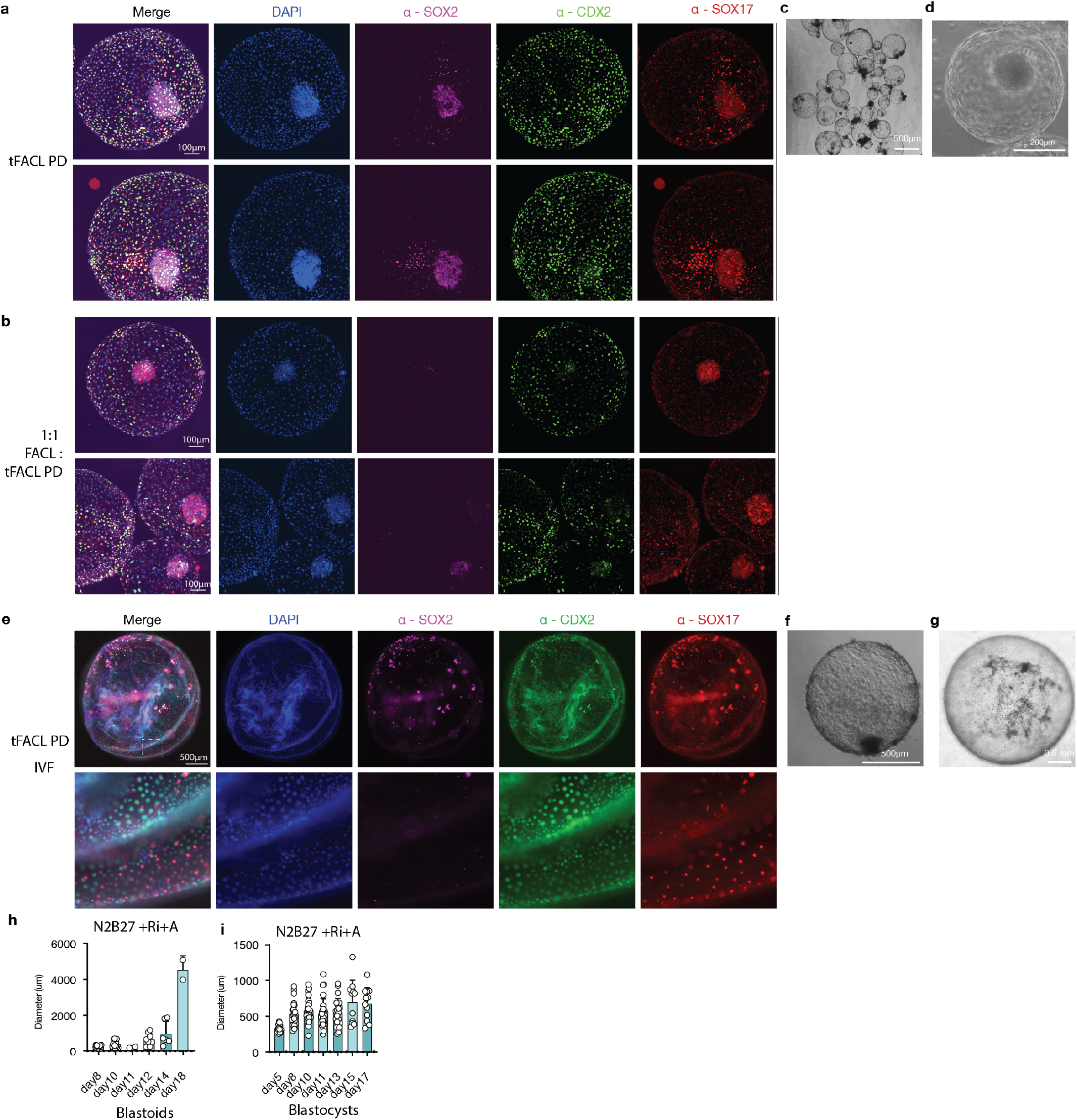
3D in vitro growth culture immunofluorescence staining. Immunostaining of bovine blastoids grown in the ClinoStar incubator at day 16 for epiblast marker SOX2 (magenta), hypoblast marker SOX17(red) and trophectoderm marker CDX2(green) in **a**. tFACL+PD media. **b**. A 1 to 1 mix of FACL and tFACL+PD. **c-d**. Phase-contrast image of bovine blastoids grown in the ClinoStar incubator. **e**. Bovine IVF blastocyst grown in in the ClinoStar incubator at day 16 for stained as in a-b. **f-g**. Phase-contrast image of in vitro grown bovine blastocyst. **h-i**. Quantification of invitro grown blastoids and blastocysts on N2NB27 with rock inhibitor (Y27632) and activin A as reported in ^34^.

**Extended Data figure 5.**
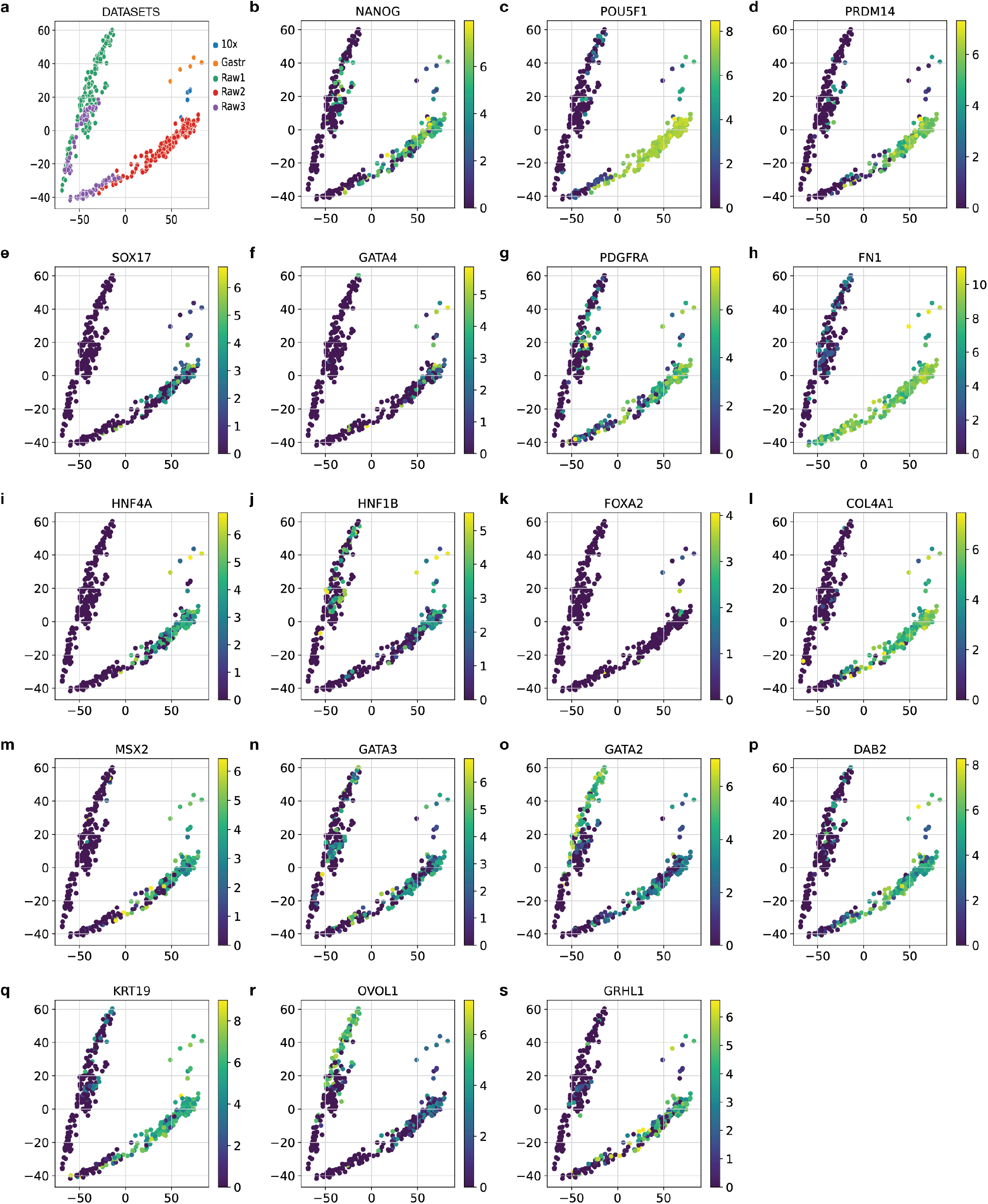
Principal component analysis (PCA) heatmaps of pseudo bulk conversion of blastoid data. **a**. Color by dataset. Epiblast markers: **b**. NANOG. **c**. POU5F1(OCT4). **d**. PRDM14. Hypoblast markers: **e**. SOX17. **f**. GATA4. **g**. PDGFRA. **h**. FN1. **i**. HNF4A. **j**. HNF1B. **k**. FOXA2. **i**. COL4A1. **m**. MSX2. Trophectoderm markers: **n**. GATA3. **m**. GATA2. **p**. DAB2. **q**. KRT19. **r**. OVOL1 **s**. GRHL1.

**Extended Data figure 6.**
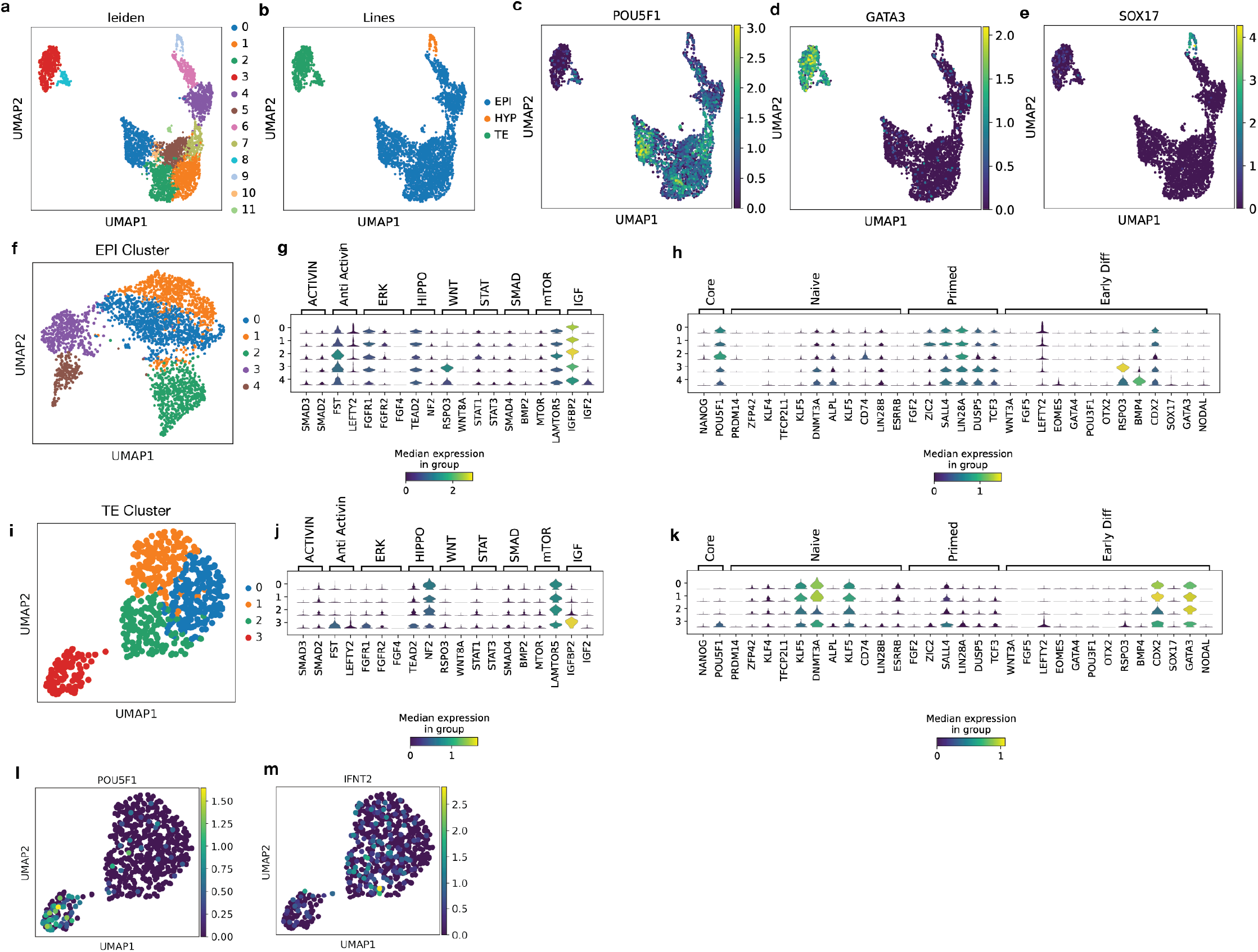
Blastoid TS and ES sub clustering analysis. **a**. UMAP of blastoid data and clustering analysis. **b**. Cluster allocation. **c**. Heatmap of epiblast marker POU5F1(OCT4). **d**. Heatmap of trophectoderm marker (GATA3). **e**. Heatmap of hypoblast marker SOX17. **f**. UMAP of epiblast subclusters **g**. Violin plot comparison of different signaling markers. **h**. Violin plot comparison of different pluripotency markers. **i**. UMAP of trophectoderm subclusters **g**. Violin plot comparison of different signaling markers. **h**. Violin plot comparison of different pluripotency markers. **l**. Heatmap of epiblast marker POU5F1(OCT4) within the TSC subcluster indicating an early blastocyst like subpopulation. **m**. Heatmap of INFt transcript INFT2 expression within TSC subcluster.

**Extended Data figure 7.**
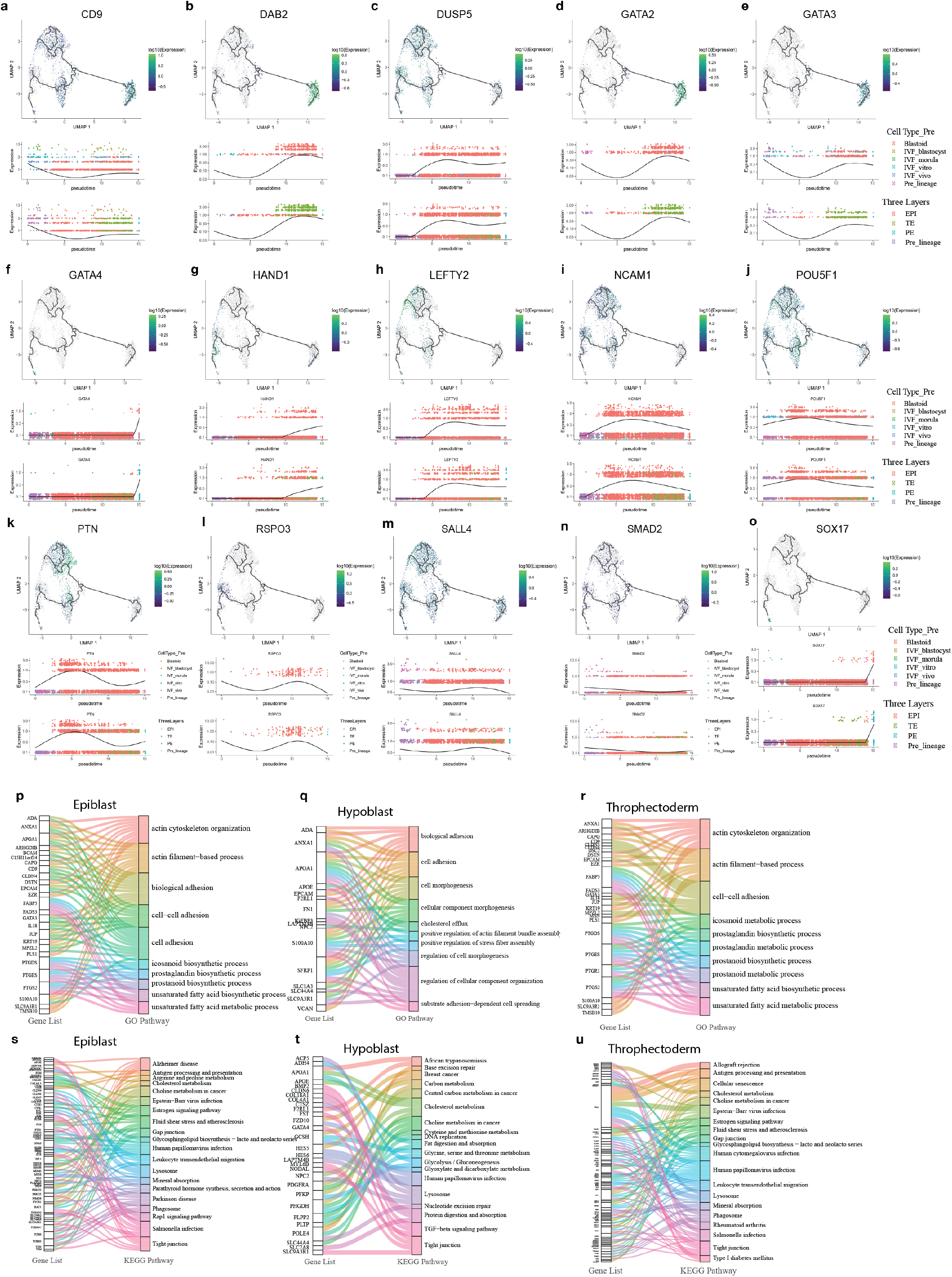
RNA velocity and pathway analysis. **a-o**. Expression heatmap and pseudotime analysis of different markers. **p-r**. Alluvial diagram of Go pathways of differentially expressed genes (DEG) in each cluster **s-u**. Alluvial diagram of KEGG pathway of DEGs.

**Extended Data figure 8.**
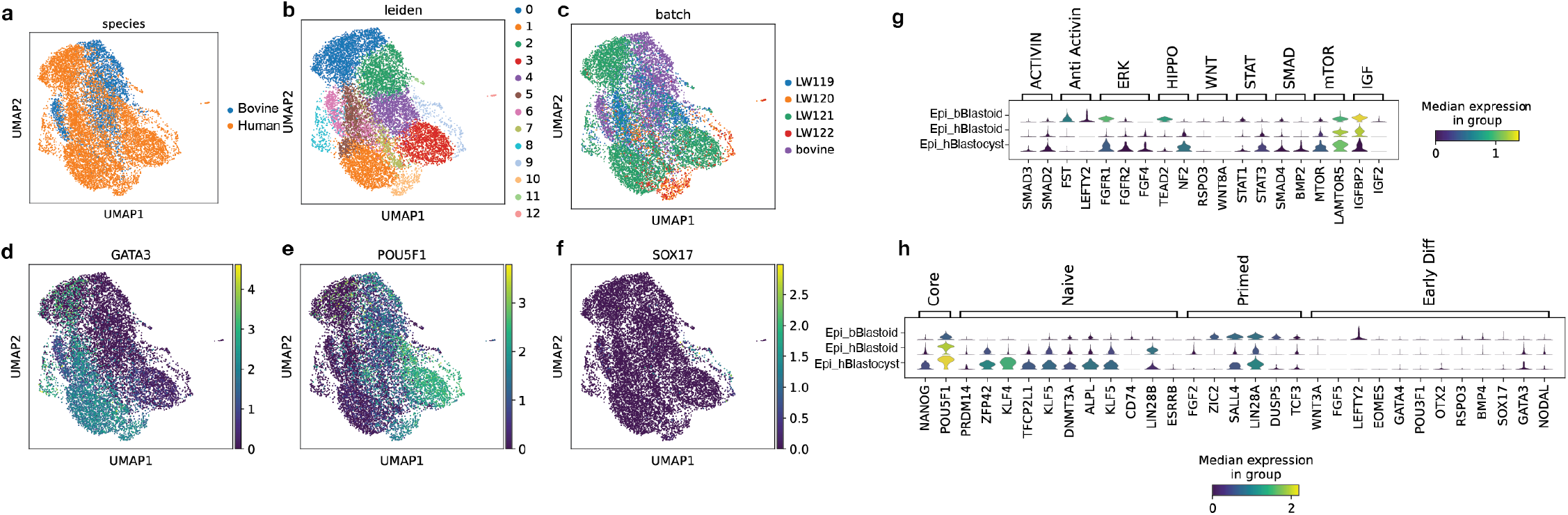
Human blastoid, blastocyst and Bovine blastoid scRNA-seq comparison. **a-c**. UMAP of data integration. **d**. Heatmap of trophectoderm marker (GATA3). **e**. Heatmap of epiblast marker POU5F1(OCT4). **f**. Heatmap of hypoblast marker SOX17. **g**. Violin plot comparison of different signaling markers. **h**. Violin plot comparison of different pluripotency markers.

